# Rethinking success, integrity, and culture in research (part 1) — A multi-actor qualitative study on success in science

**DOI:** 10.1101/2020.02.12.945733

**Authors:** Noémie Aubert Bonn, Wim Pinxten

## Abstract

**ABSTRACT:** *Background:* Success shapes the life and careers of scientists. But success in science is difficult to define, let alone to translate in indicators that can be used for assessment. In the past few years, several groups expressed their dissatisfaction with the indicators currently used for assessing researchers. But given the lack of agreement on what should constitute success in science, most propositions remain unanswered. This paper aims to complement our understanding of success in science and to document areas of tension and conflict in research assessments.

*Methods:* We conducted semi-structured interviews and focus groups with policy makers, funders, institution leaders, editors or publishers, research integrity office members, research integrity community members, laboratory technicians, researchers, research students, and former-researchers who changed career to inquire on the topics of success, integrity, and responsibilities in science. We used the Flemish biomedical landscape as a baseline to be able to grasp the views of interacting and complementary actors in a system setting.

*Results:* Given the breadth of our results, we divided our findings in a two-paper series, with the current paper focusing on what defines and determines success in science. Respondents depicted success as a multi-factorial, context-dependent, and mutable factor. Success appeared to be an interaction between characteristics from the researcher (Who), research outputs (What), processes (How), and luck. Interviewees noted that current research assessments overvalued outputs but largely ignored the processes deemed essential for research quality and integrity. Interviewees sustained that we need a diversity of indicators to allow a balanced and diverse view of success; that assessments should not blindly depend on metrics but also value human input; that we must value quality over quantity; and that any indicators used must be transparent, robust, and valid.

*Conclusions:* The objective of research assessments may be to encourage good researchers, to benefit society, or simply to advance science. Yet we show that current assessments fall short on each of these objectives. Open and transparent inter-actor dialogue is needed to understand what research assessments aim for and how they can best achieve their objective.

*Trial Registration:* osf.io/33v3m

## BACKGROUND

Excellence is a prominent theme in any funding scheme, university mission, and research policy. The concept of excellence, however, is not self-explanatory. Apart from the fact that excellence is hard to define, it is complicated to translate it into concrete criteria for evaluating whether researchers are successful or not in their pursuit of scientific excellence. Nonetheless, in today’s highly competitive setting where talent is plenty and money is tight, determining evaluation and assessment criteria is a necessity.

When researchers are being assessed, it is important that the criteria used for determining success are compatible with our concepts of scientific excellence. However, with poorly defined concepts of excellence (e.g., 1) and assessment criteria that raise considerable controversy, there is no guarantee that this is actually the case.

The issue has increasingly attracted the attention of influential voices and fora, which resulted in a growing number of statements and documents on the topic, including the Declaration on Research Assessment (DORA; 2), the Leiden Manifesto (3), The Metric Tide (4), and more recently the Hong Kong Principles for Assessing Researchers (5). In a review of 22 of these documents, Moher and colleagues pointed out that current research assessments are open for improvement, particularly in further addressing the societal value of research, in developing reliable and responsible indicators, in valuing complete, transparent, and accessible reporting of research results as well as reproducibility, and in providing room for intellectual risk taking (6). As many of the documents mention, however, changing scientific assessment is not straightforward and is likely to face resistance from diverse parties. One of the reasons for this resistance may be the complex inter-actor exchange that governs research and academia. As the European Universities Association (EUA) made clear in a recent report, research institutions, funders, and policy makers must “work together to develop and implement more accurate, transparent and responsible approaches to research evaluations” (7, p. 13). But although certain actors such as researchers and scientific editors have been highly involved in the debate, other actors have been largely missing from the discussion.

The present research contributes to this discussion by expanding the understanding of success in science and by unraveling the connections between success and research integrity. We use the Flemish biomedical research landscape as a lens to study what success means in science, how it is pursued, and how it is assessed. Noticing that most research on research integrity captures the perspective of researchers and research student (8), we decided to extend our understanding of success and integrity by also capturing the views of policy makers, funders, institution leaders, editors or publishers, research integrity office members, research integrity network members, laboratory technicians, and former researchers who changed career. Our findings, divided in a two-paper series (see 9 for our associate paper describing the problems that currently affect academia), our findings resonate with past efforts by suggesting that, in their current state, research assessments may fuel detrimental research practice and damage the integrity of science. In this first paper, we discuss the way in which different research actors perceive success in science.

## METHODS

### Participants

The present paper reports findings from a series of qualitative interviews and focus groups we conducted with different research actors. This qualitative work was part of the broader project Re-SInC (rethinking success, integrity, and culture in science; the initial workplan is available at our preregistration (10)).

In Re-SInC, we captured the views of different research actors on scientific success, problems in science, and responsibilities for integrity. Being aware that the term ‘research actor’ may be ambiguous, we defined research actors as any person having a role in the setup, funding, execution, organisation, evaluation, and/or publication of research. In other words, we included actors linked to the policing, the funding, the evaluation, the regulation, the publishing, the production (i.e., undertaking the research itself), and the practical work of research, but we did not include sole consumers of science or end users of new technologies.

We used Flanders as a setting, including participants who either participate in, influence, or reflect (directly or indirectly) upon the Flemish research scene. In most cases, the interviewer did not know the participants before conducting the interviews and focus groups. In selecting participants, we aimed to capture the breadth of the Flemish research scene. Using Flanders as a research setting had the advantage of allowing us to capture perspectives from an entire research system in a feasible setting. The Flemish research scene comprises of five main universities and a number of external research institutes, major funding agencies, a federal research policy department, and one advice integrity office external to research institutions. We chose to concentrate our research on three of the five universities, and to include partnering European funding and policy organisations as well as international journals and publisher to build a realistic system sample. When participants were affiliated with a university, we focused on the faculty of biomedical sciences. Given the exploratory and qualitative nature of this project, we did not aim for an exhaustive nor a fully representative sample. Our objective was to shift the focus from the narrow view targeting mainly researchers to a broader view that includes a broad range of research actors. Accordingly, we maximized the diversity of participants in each actor group to ensure that each group encompassed a wide range of potentially different perspectives.

Our main actor categories are PhD students, post-doctoral researchers (PostDoc), faculty researchers (Researchers), laboratory technicians (LT), policy makers and influencer (PMI), funding agencies (FA), research institution leaders (RIL), research integrity office members (RIO), editors and publishers (EP), research integrity network members (RIN), and researchers who changed career (RCC). The composition of each actor group is detailed in Table 1.

**Table 1.**
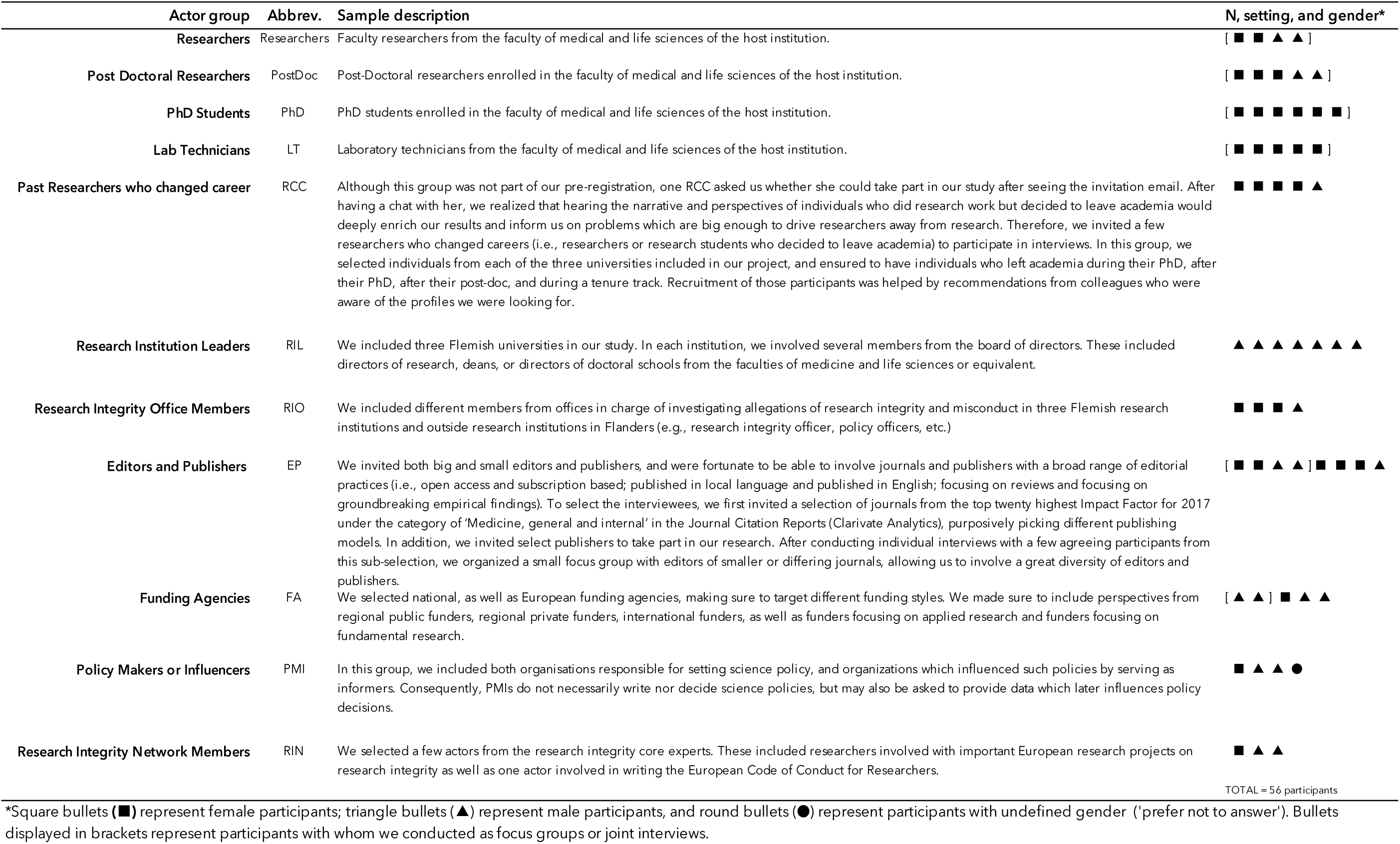
Demographics of participants.

It is important to keep in mind that the research world is complex and not organized in distinct actor groups. Consequently, participants could often fit in more than one category, and sometimes felt the need to justify circumstances that would make them fit in the category we selected. Before the interview, we asked participants whether they agreed with the category we assigned them in, and we refined and exemplified the definitions of our actor groups to reflect the participants’ distinctions (i.e., further explaining the slight differences between the groups planned in the registration and those used here).

### Recruitment

We used several recruitment strategies. For the focus groups with PhD students and researchers, we circulated an email to everyone in the faculty of biomedical and life sciences of the host university and invited them to register on an interest list. We later scheduled a convenient time with those who registered. We used a similar strategy for the focus group of editors and publishers, but circulated the invitation in a conference of scientific editors. For focus groups with lab technicians and postdocs, key players helped us recruit and organize the focus group.

For interviews, we invited participants directly via email. We sent up to three reminder emails, but did not pursue further if no response was obtained at the third reminder email. All participation was on a voluntary basis.

### Design and setting

We conducted semi-structured interviews and focus groups, meaning that we asked broad questions in an open manner to target main themes rather than specific answers. All interviews and focus groups were audio recorded and transcribed verbatim. Details about the tools used to guide the interviews and focus groups are available in the tool description below.

To maximise transparency, we provide an extended descriptions of the interviewer and the setting of the interviews in Appendix 1 and a copy of the COnsolidated criteria for REporting Qualitative research checklist (COREQ) in Appendix 2.

### Ethics and confidentiality

The project was approved by the Medical Ethics Committee of the Faculty of Medicine and Life Sciences of Hasselt University (protocol number CME2016/679), and all participants provided written consent for participation and for publication and dissemination of the findings from this project. A copy of the consent forms is available in the registration of this project (11). We protected the confidentiality of participants by removing identifiers from quotes included in the text. Nonetheless, Flanders is a small research system and given our actor-specific sample, personal identification within quotes remains a risk despite our efforts. To further protect participants’ confidentiality and avoid that identification of individual quotes lead to identification of all quotes from the same participant, we decided not to specify respondents in individual quotes, but to refer only to actor groups.

Following this reasoning, we are unable to share full transcripts, but attempted to be as transparent as possible by providing numerous quotes in the text, in tables, and in appendices.

### Tool

To build our focus group guide, we inspired our style and questions from the focus group guide developed by Raymond De Vries, Melissa S. Anderson, and Brian C. Martinson and used in a study funded by the NIH (12). We obtained a copy of the guide after approval from the original authors, and revised the guide to tailor questions to the topics we wished to target, namely ‘success in science’ and ‘responsibilities for research integrity’. We revised our focus group guide several times before data collection and discussed it with Raymond De Vries — expert in qualitative inquiries and part of the team that built the original guide upon which we inspired ours. We built interview guides based on our revised focus group guide. We adapted specific questions (e.g., responsibilities, evaluation) to each actor group, but preserved the general structure and themes for all interviewees. A general version of the interview and focus group guides are available in Appendix 3 and 4. More specific group guides can be provided upon request. All guides were constructed around the following four topics:

i. **Success in science**: which characteristics are most important to advance a researcher’s career? What leads to success? What are indicators for success?
ii. **Current problems** (including misconduct and questionable research practices): Do you have experience with research that crossed the lines of good science? How can we draw the line, what are red flags? Why do bad practices happen? Can they happen to anyone?
iii. **Responsibilities towards integrity**: What is your responsibility towards integrity? Where does it end? Who else is responsible? In what ways are other actors responsible?
iv. If you were granted a fairy wish and could **change one thing in how science works**, what would you pick?

It is important to consider that the interview guide was not used mechanically like a fixed questionnaire, but sometimes shortened or expended to capture responses, interest, and to respect time constraints.

### Analysis

Recordings were first transcribed verbatim and, where necessary, personal or highly identifiable information was anonymized. We analyzed the transcripts using an inductive thematic analysis with the help of NVivo 12 Software to manage the data. The analysis proceeded in the following order:

i. **Initial inductive coding**: NAB first analyzed two focus groups (i.e., researchers and PhD student) and five interviews (i.e., RIL, RIO, PMI, RCC, and RIN) to have an initial structure of the themes targeted. In this step, she used an inductive thematic analysis (13) while keeping the three main categories — i.e., success, integrity, and responsibilities — as a baseline. Using the inductive method allowed us not to limit our analysis to the order and specific questions included in guide, but to also identify and note themes that were raised spontaneously or beyond our initial focus.
ii. **Axial coding**: With this first structure, NAB and WP met and took a joint outlook at these initial themes to reorganize them in broader categories and identify relationships between categories. For this step, NAB built figures representing connections between the main themes, and refined the figures and the codes after the meeting.
iii. **Continued semi-inductive coding**: NAB continued the coding for the remaining transcripts, sometimes coding deductively from the themes already defined in steps 1 and 2, and sometimes inductively adding or refining themes that were missing or imprecise.
iv. **Constant comparison process**: NAB and WP repeated the axial coding and refining several times throughout this process, constantly revisiting nodes (i.e., individually coded themes) by re-reading quotes. The nodes and structure were then discussed with RDV to reconsider the general organisations of the nodes. This constant comparison process is common in qualitative analysis, and is commonly used, for example, in the Qualitative Analysis Guide of Leuven (QUAGOL; 14). This repeated comparison led to a substantially solid set of nodes which later guided further coding in a more deductive manner, though we made efforts to remain open to possible new themes in respect of our inductive analysis.
v. **Lexical optimization**: Finally, after having coded all transcripts, NAB and WP further discussed the choice of words for each node and reorganized the themes to ensure they have an ideal fit with the data they are describing. NAB and RDV met to have a final outlook of the general structure, and to reorganise the nodes in clean and natural categories.

### Limitations to consider

A few points are important to consider when interpreting our findings. First, given the exploratory and qualitative nature of this project, our sample is neither exhaustive nor fully representative. We chose to ask for personal perspectives rather than official institution or organisation views since we believed it would allow us to capture genuine beliefs and opinions and to avoid rote answers. We thus encouraged participants to share their personal thoughts rather than the thoughts that could be attributed to their entire actor groups, institution, or organisation. We consider that these personal beliefs and opinions are crucial in shaping the more general views of organisations, yet we urge our readers to remain careful when making group comparison and generalisations.

Adding to the above concern, it is important to keep in mind that the research world is complex and not organized in distinct actor groups. Participants could often fit in more than one category by endorsing several research roles. As we mention above, we asked all participants whether they agreed with the category we assigned them in, and we refined and exemplified the definitions of our actor groups to reflect the participants’ distinctions. Yet, we must consider each actor category not as a closed box with characteristic opinions, but as a continuum which may or may not hold divergent views from other actor groups. Our findings help capture views which may have been overlooked in past research which focused on researchers, but should not be used to discriminate or represent the opinions of entire actor groups.

Finally, it is important to consider that given the richness of the information gathered, certain findings may be displayed with greater importance than others simply based on the authors’ personal interests. We were careful to include also the views we disagreed with or found to be of limited interest, yet it is inevitable that some of the selection and interpretation of our findings was influenced by our own perspectives. To maximise transparency on the genuine views of our informers, we supplement our interpretation of the findings with quotes whenever possible.

## RESULTS

The purpose of this paper is to retell, connect, and extend on the issues that the different actors raised in our study. Aiming to maximise transparency and to minimise selective reporting, we provide numerous quotes and personal stories to illustrate our claims. The result, however, is a lengthy paper in which we explain the breadth of the concerns raised by our participants. Given the length of the resulting paper, a short summary of results is available at the end of the results section, and select findings are re-examined and extended in the discussion.

### Researchers’ personal successes

Before reporting on the views of all interviewees on research success, we believed it would be important to look at the answers of researchers and research students. Focus groups with researchers and research students comprised an additional question in which we asked participants to describe their personal satisfactions and successes. Given the limited number of researchers and research students involved in our research, it would be naive to infer that our findings represent the breadth of researchers’ view on success. Nevertheless, we believed that capturing what researchers and research students describe as ‘satisfying’ was important to understand and contextualise the general perspectives of success in science.

In their answers, interviewees described a number of factors which made them feel satisfied or which they interpreted as personal success. First, PhD students and Post-Doctoral researchers strongly supported that **making a change** in practice was something that was central to them.

> *I agree with the fact that that feeling that something is done with what you found is crucial for your own feeling. […] I think that’s crucial. Even more than the publications or the… yeah… (PostDoc)*
>
> *Yeah it was part of my motivation to give something back to the clinical field by doing research. (PhD student)*

For PhD students, realising that their results would remain theoretical or would be too small to make a difference was raised as one of the disappointment they faced in research.

> *P1: If I can help people by doing this project, that gives me a lot of satisfaction I think*.
>
> *P2: That’s true but that was also my first idea when I started, but I have to be honest, my project is so fundamental that I’m almost finishing up, and I don’t see anything that will be going to the clinic for years or something. So at that point for me it was a bit disappointing, because… Ok, I wanted to, but I’m so fundamental, basically really molecular stuff, that I don’t see it to get really…*
>
> *[…]*
>
> *P3: Yeah I think for me it’s the same. Because I’m working on a project that’s like this very tiny subset of a subset of [specialised] cells. And then at the beginning you think ‘I’m going to change the field with this research’, but yeah I don’t know. (PhD students)*

Although some researchers also supported that translating their findings in practice was satisfying, they acknowledge that **theoretical knowledge** or simply following their **curiosity** became their “*main drive*”, or at least provided its share of satisfaction.

> *“For me it’s good if it goes this direction* [i.e., is translated in practice] *but also just creating new knowledge which doesn’t really directly impact people, I think is also very very interesting, or I’m also very passionate about that. So it shouldn’t always have an implication*.*” (Researcher)*

For researchers, external satisfactions, such as **peer appreciation**, or **fulfilling institutional requirements** were seen as “*also very important*” to personal satisfaction, but as secondary aspects which were not enough for feeling completely satisfied.

> *“I also have some… still some criteria which I have to do that I also think about those things. But I don’t feel bad about it that it’s my only drive for some things that it’s just publication. On the other hand I also feel that I cannot be satisfied alone by those things*.*” (Researcher)*

Finally, researchers and research students added two intriguing dimensions to the concept of success. First, they sustained that successes are personal, and that each researcher will likely be successful in different ways. In this sense, personal success was seen to reflect aptitudes and skills in which individuals excel, rather than a universally shared idea.

> *“[In my group, we don’t have strict requirements], and I think it’s very beautiful because we have [dozens of] PhD students and they’re all — or 99% of them are — successful, but they are so different in being successful. Some are really being successful in the number of publications, some of them are really successful in the network they have with other companies, with other research institutes, some of them are really successful in the perseverance to do something really new and to make it happen, only if there’s a small study on 20 patients, but it’s so new and they will really make it happen in the hospital. So, they’re so successful on so many different levels and I really like the fact that we don’t judge them all in the same way because they can be themselves and be successful in the way that they want to be successful*.*” (PostDoc)*

The need for diversity of successes was thus valued, even though it was acknowledged to be a rare feature in research assessments. A second intriguing dimension was raised by a Post-Doctoral researcher who pointed out that, even within individual researchers, personal conceptions of success may be mutable, likely influenced by career stages, work environments, and expectations of others.

> *“P1: For me I think my idea on what success is* ***changing a lot*** *of course. When you’re a PhD student you just want a breakthrough in your project, that’s success, and then by the time you’re finishing your PhD you’re looking at what… Is there a possibility for post doc then you realise ‘OK they’re counting publications, they’re doing this’ and then you’re looking around and then you sometimes get this mixed feeling of someone who you feel was not very creative or did not have to do a lot of work themselves, it was very guided and clear steps, and they have a lot of publications and so they get a post doc position. And then that’s sometimes difficult, and you think like ‘How does this work here?’ […] then I went into more research coordination and then I was in a [different group] and then it was all the time about metrics. Because the money was divided by metrics, and it was like publications and project funding and… And then I felt like everything revolved around that. It wasn’t important anymore like what projects we’re doing as long as it was a project on that funding channel because that counted higher on the metrics and… So ok, and* ***then you’re really like that***. *And now being here in this setting I’m really seeing the impact of research*. ***Now it’s changed again***. *Now it’s really like that kind of research where you can make a difference for an organisation, for patients… That’s the thing that’s success. And I think that maybe like you say that in the long run that’s what you have to do. But it’s kind of the short-term mechanisms, and not always…*
>
> *P2: R: Yeah, I think that the definition of success is highly dependent of the institute and the environment you’re in like you’re mentioning. And if you’re constantly told ‘This is how we measure success’ then…*
>
> *P1:Yeah, so* ***then you’re really guiding yourself to get those key indicators***.*” (PostDoc, bold added for emphasis)*

In other words, interviewees revealed that personal success was a mutable variable which appeared to change depending on contexts, demands, and career stages.

### Inter-actor views on success

Now that we have glanced at the perspectives of researchers and research students, let’s look at the views of all research actors on the more general idea of success. In order to avoid rote answers, we asked about success indirectly. Rather than asking ‘What is success in science?’, we asked interviewees about ‘What makes researchers successful?’.

In their answers, interviewees mentioned several factors which they believe are essential or useful in becoming a successful researcher. We classified these factors in four main categories: factors visible in the *researchers* themselves (Who), factors from the research *process* (How), factors from the research outputs (What) and, unexpectedly, factors related to *luck*, which was thought to play an important role in success^1^. Figure 1 illustrates the different categories we captured.

**Figure 1.**
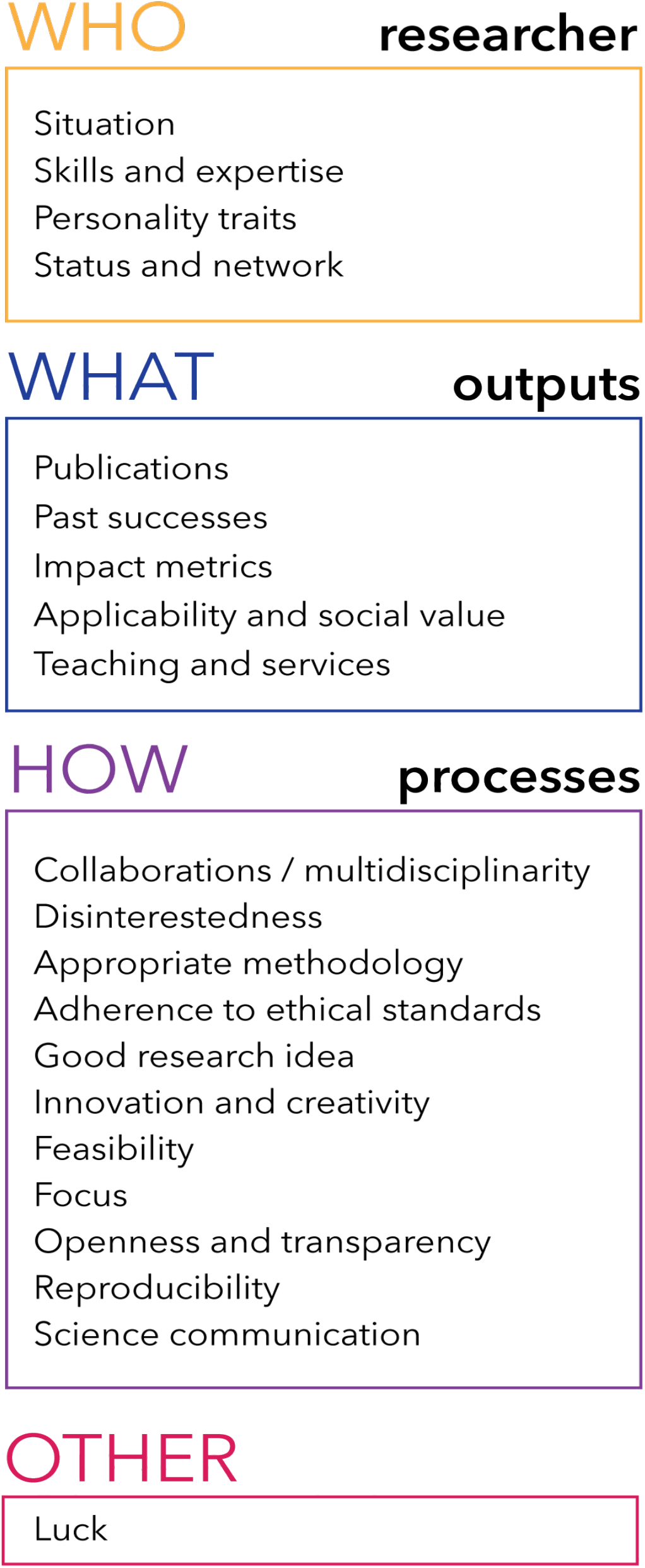
Main themes captured as determinants of success in science.

#### Who

##### Researcher

A number of features which are related to the *researchers* themselves were considered important in determining and yielding success. While all these individual factors were said to play a role in producing success, they were also described as indicators to look for when selecting researchers for a position and thus influenced careers and promotions. Among those, participants highlighted *personal traits*, such as ambition, passion, rigorousness, and intelligence, as well as acquired *skills and expertise*, such as business potential, management skills, writing skills, and scientific expertise. Certain respondents also believed that success could be influenced by specific *situation* in which individual find themselves. In this regard, gender and ethnicity were mentioned as possible obstacles — through pregnancy leaves, family obligations, prejudice, or language inequalities — or advantage for success — through employment quotas. Along the same line, childlessness and celibacy were mentioned as advantages for yielding success since they allowed researchers to devote more time to their work.

Beyond the advantage that extra time and flexibility could provide, it was seen by some as a condition to a successful research career. Some interviewees sustained that researchers and research students *should* be able to devote themselves to their career by being mobile and by working beyond regular schedules and conditions.

> *“I think people have to realize when you do a PhD, it’s a stressful thing, you really are going to get the highest degree there is at a university, it doesn’t fit between 9 and 5*.*” (RIL)*
>
> *“…being passionate about science is almost like being an artist. You live in poverty because you want to pursue your art*.*” (PMI)*.
>
> *“That’s also what we ask for, excellence for people when they come here. […] Usually those people need to have been abroad for at least six months. But if it is two years it’s better. So these are important factors to create excellence*.*” (RIL)*

Many of the researchers who changed career mentioned that the expectation that they should sacrifice family life and private comfort for science played a role in their decision to leave academia. We will explore this idea further when discussing unrealistic expectations in the associate paper (9). Finally, the network and status that researchers bring along with them was also seen as determinant to success. Having an established network and personal recognition from peers was thought to be key to success.

#### What

##### Outputs

Indicators which provide information about what researchers have accomplished were univocally considered crucial in determining success. Among those, high academic grades, past success in obtaining funding, publications, and publication metrics (e.g., impact factor, citations, H index) were mentioned as currently being used for determining success, although not all interviewees agreed on the individual value of these determinants. In addition, less traditional products of research were also mentioned, such as the applicability and societal value of the research findings and the researcher’s involvement in teaching and services (i.e., mostly referred to as serving on institutional boards, committees, and scientific societies).

#### How

##### Processes

But features which indicate ‘how researchers work’ (i.e., *processes*) were also deemed integral to success, regardless of the output they generate. On the one hand, some processes were thought to play a part in the success of individual research projects. Collaborations, multidisciplinarity, appropriate methodology, adherence to ethical requirements, good and innovative research ideas, feasibility, and focus were all viewed as pathways to achieve good outputs and related successes. On the other hand, respondents also identified a number of processes which they considered impacted beyond individual projects and were essential to the success of science at large. Openness and transparency, for example, were repeatedly viewed as important aspects of the collegiality which promotes the success of science as a common goal. One interviewee explained that openness was “*very important to help the research enterprise because it’s really about facilitating the fact that other people can build upon a research”* (EP). Along the same lines, reproducibility was considered as the “*most important thing*” (RIL) and as “*a very important element in science*” (PMI). Yet, interviewees noted that reproducibility is “*often lacking*” from research (PMI), and that replication studies are under-appreciated in current success assessments (Researcher) or even possibly wasting research money (RIL). Finally, public engagement, mainly in the form of communicating scientific findings to the public, was also mentioned as part of the broader scientific success by building trust in science and by potentially contributing to the quality of research.

What truly differentiated *outputs* from *processes* was the perspective that the latter contributed to success regardless of the final result.

#### Other

##### Luck

Interviewees also attributed success to *luck*, a feature which transcended outputs, processes, and individuals. In our analysis, we discerned three different meanings of what it meant to be ‘lucky’ in science. First, researchers could be considered lucky if they worked with distinguished colleagues or in established labs, given that such settings maximized the opportunities for obtaining high end material, publications, and grants. This first meaning brings back the idea of the network that researchers bring with them, and adds an element of arbitrariness to the control that researchers have in building their network. Second, luck was also employed to refer to unexpected evolutions and trends, such as working on a topic which suddenly boomed in visibility and media attention or being “*somewhere at the right moment at the right time*” (FA). In this second signification, luck was perceived as something that one could partially create, or at least grasp and maximize. Finally, luck was sometimes attributed to the output of research results, with positive findings being lucky, and negative findings being unlucky. In this last sense, luck was a factor that was out of researchers’ control and independent of their skills. In all three senses, luck was both described as something that had helped mediocre researchers move ahead in their career and as something that had wrecked the success of otherwise talented researchers.

### Current assessments of success

In the above section, we describe the different features which were used to describe success in science. Although this broad array of features described the overall picture of success our interviewees revealed, current research assessments do not necessarily value these elements equally. In answering our question about ‘What makes researchers successful?’, several interviewees spontaneously identified the tension between what currently determines success — through formal rewards — and what they believed *should* determine success in science.

> *“Hm… What the current situation is, or what I think success should be? (laughs)” (EP)*
>
> *“I think that you have different views on looking on it. You have the measurable parts, and you have the non-measurable part. And I think that these two are sometimes in contradiction*.*” (RIO)*
>
> *“…I started this PhD project because I wanted to have results useful to clinical practice, and I said “I want to do this”. And [my supervisors] were already saying for a year “No, no, it’s not interesting, no we shouldn’t do that*.*” and I said “I want to do this, or my project failed for me*.*” […] Ok, I know it’s not going to be so big that it’s so interesting for journals, but I think for our clinical field, for Flanders, it’s important that we do a study like that. And it was… that was the chapter that people from clinical practice were most interested in too. So… I think when you ask us ‘What is success in research’, we’ve got our own points of success, and what we know that’s expected from us by the system. So those are two different lists. (laughs)” (PhD)*

More precisely, assessments of success were described as currently focusing on research outputs — generally measured through rigid metrics and quantity indicators — while largely ignoring other important features from the process through which science is performed. In this respect, several interviewees mentioned that although output-related successes helped researchers advance their career and made them feel satisfied in some way, they also felt a lack of reward for processes which provided an indication about the quality of the work, its usability for the scientific community, and the quality of science as such.

Despite a general agreement on the need to reintroduce processes in research assessments, respondents sometimes contradicted each other when we asked them to give precise examples of outputs that bothered them, or to describe processes they thought should be valued more in research assessments.

#### Disagreements on outputs

##### Publications

The emphasis on publication in research assessments raised such a disagreement. Some respondents considered that relying on publications to measure success was problematic and even damaging for science, while others saw publications as a necessary and representative measure of scientific success. We distinguished three arguments against, and three arguments in favor of using publications as the main indicator for assessing research success (Figure 2). Illustrative quotes are available in Appendix 5.

The first argument against using publications as a main indicator of success was based on the idea that publications, constitute a *reductionistic* measure of success. In other words, using publications as the main measure for success ignored “*other* very *important contributions to the scientific enterprise*” (EP). Additionally, the reductionistic scope of publications was said to sometimes unjustly penalised researchers who “*have the qualifications to be good researcher*” (PhD student) but are simply unsuccessful in publishing their results. The second argument against focusing on publication for evaluating success resided on the belief that publications are an *arbitrary* measure which does not represent merit, efforts, and quality. Researchers and research students in particular worried that publications often resulted from arranged connections rather than from high scientific value or efforts, an argument we will discuss further in the associated paper (9). Researchers and students also supported that high impact publications in recognized journals were not necessarily of high quality, making the link between papers and quality arbitrary. Adding to this, several interviewees supported that publications could be a mere matter of luck. As a third argument against focusing on publications to evaluate researchers, interviewees worried that the increasing dependency of researchers on their publication output (i.e., the publish or perish culture) may introduce *perverse* incentives which might threaten the integrity of research. On the one hand, publication pressures may tempt researchers to engage in questionable practices to maximise their publication output. On the other hand, many interviewees also noted that the emphasis on publications sometimes shifted the main objective of research projects towards ‘sexy’ and ‘publishable’ topics rather than topics that are ‘interesting’ or ‘relevant’ to advance science.

Despite these three arguments against the focus on publications for evaluating success, a number of respondents also identified arguments in favour of such focus, sometimes directly opposing the arguments introduced above. First, publications were described as *necessary* aspect of scientific advancement, and the emphasis that evaluations give to publications was seen as a way to ensure that researchers keep publications in the forefront of their priorities. Second, some respondents described publications as *representative* indicators good research, and merit. In fact, considering that publications are the endpoint of an extensive and difficult process which could not happen without hard work, some argued that publication outputs helped identify good researchers. Finally, publications were also described by many as the only tangible way to value and *measure* science, which added to the credibility of research assessments.

##### Impact factor

Similar conflicts were observed when looking at the impact factor. First, the impact factor was acknowledged by many as a being useful for its *measurability*, its *simplicity*, its *acceptability*, and even — as some mentioned — for its perceived correlation with the *quality* of the review process.

> *“So if you hire a PhD student, but even more if you hire a postdoc or a young professor, then they evaluate it of course. And then, the bibliometric parameters are much more important so you look at the number and the quality of publications*. ***How do you measure the quality, of course the impact factor***. *So if somebody with a Nature paper comes of course this person is considered to be more ‘valuable’, in quotation marks, and gets a higher score in the end and probably the job, compared to a person with a publication record which has lower impact factors. So impact factors are still very important, and grants… (RIL, bold added for emphasis)*
>
> *Of course what you always want to have is one of the two champions that are really picky in the graph, but I think for us it’s also important to really see that the whole group is evolving to improved* ***quality as measured by the impact factor*** *and of course I know the discussion that this is only one way to look at quality, but* ***it’s still the most accepted way to look at quality*** *I think, in our field. (RIL)*
>
> *I have to say that generally* ***there is a big correlation between the impact factor and the quality*** *of the content… (EP, bold added for emphasis)*
>
> *OK, when we select something, somebody for an academic position, we will look at publications, at the numbers, and below 10 you will never get something in an academic position, below 10 papers. Of course, suppose somebody comes with two Nature, one Lancet, and one NEJM, then we have to re-think. So… In a way, today it’s still a balance between numbers and impact factors, it’s still playing a role. But the whole issue is that there is something which goes together*. ***A journal with a high impact factor has to improve its review process***. *Because you cannot keep your high impact… I think that when you send your paper to Lancet or NEJM, you will have tough review. While when you send it to a low impact factor journal, […] you can send a completely fake paper to reviewers who will judge it perfect and let it publish. (RIL, bold added for emphasis)*

Nonetheless, using the impact factor as a measure of success yielded overwhelmingly negative responses. Most participants mentioned that the impact factor was not adapted to their disciplines, not representative of the impact of individual papers, open to biases and manipulation, and even disrupting science by discouraging research in fields with traditionally lower impact factor.

> *I think [current metrics are]* ***far too simple***. *You know like impact factor is* ***useless I think in evaluating the importance of an individual paper***, *because impact factor relates to a journal. So it’s not an article level measure of any kind. (EP, bold added for emphasis)*
>
> *Publishing is important but I hate the impact factor thing. I would more look into the quartile thing, if you are in a* ***field*** *that has low impact factors but you are in the top ten of your field, that’s just fine. I mean it doesn’t have to be Nature, it can also be [a small specific journal], if that’s your top, in your field. So I think there is a tendency towards going that way but I like that a lot more than the impact factor shizzle, yuck! (RCC, bold added for emphasis)*
>
> [Interviewer] *Which [indicators] do you think are the most toxic and less representative of quality?*
>
> [Participant] *The urge to publish in Q1. […] I understand that there needs to be an impact factor, but the whole issue of the weight of an IF in the personal career of a researcher… because then I would advise anybody who wants to go in research “Please go in cancer research”. Try to get to Lancet cancer or whatever other journal, of NEJM and then you’re safe. Don’t do anything like plastic surgery or [smaller topics]… So* ***that’s one of the most toxic factors I think***. *The pressure of… Because the* ***impact factor is not reflecting really the importance of the research***. *You could say that cancer is of course important, and then you see that for instance [the biggest journal in cancer research […] they manipulate the impact factor. Of course, because when you, as an author, you don’t have enough references referring to their own journal you get from the reviewer report that you need to put those in… (RIL, bold added for emphasis)*
>
> *Well the problem with the impact factor as a standard, most appreciated metrics, even though we don’t want to do that [laughs], is that it is* ***not essentially an indicator of quality neither of the article, neither of the journal***, *but why? Because there could be less articles of lesser quality, published by renowned scientists in higher impact factor journals, and you can have a good research from scientists coming from some small country and who is not so famous internationally, and he will not, or she will not be able to publish in the higher impact factor journals because they are usually biased, and I know because I come from a country, when you read someone’s last name you usually… they can know that you are from that country [laughs] (EP, bold added for emphasis)*

A few interviewees proposed that direct citations would be more relevant in personal impact assessments, but they also acknowledged that determining the impact of individual articles using direct citations could take years, if not decades in some disciplines. Furthermore, researchers added that their most cited paper was not necessarily the one they considered most important, and that citation counts tended to refer to novelty and timing rather than to quality. Consequently, despite an overwhelming aversion towards using impact factors for scientific evaluation, concrete alternatives were more difficult to nail down.

#### Disagreements on processes

##### Science communication

The importance of science communication also raised conflicting opinions among interviewees. Many supported that sharing science through popular channels such as Twitter, youtube, Facebook, or Wikipedia should be considered in career evaluations. For example, one respondent in the PMI group raised that “*Researchers who do a lot of work on Wikipedia are not rewarded for it, but they’re doing a lot of good work!*”. The same interviewee however, later warned that appearing in the media was different than actively making the effort to communicate science, “*because then the media decides who is successful*” and “*a lot of researchers will also be successful and you will never hear of it*”. A few researchers mentioned that science communication was essential to maximize the interdisciplinary impact of one’s work, and that presenting findings in broad conferences and participating on Twitter could foster this interdisciplinarity. But some respondents also sustained that science communication and the ability to simplify and share one’s findings with different stakeholders went beyond the duty of sharing and was actually key to the core quality of the work.

> *“I mean we work a lot on, or we try to promote everything which has to do with public engagement and science communication, all these things, but if you’re not able to explain to lay people what your research is about… [moves head meaning it’s not a good sign]. I think it’s a sort of, how do you call it, a litmus test in a certain way […] Sometimes sort of public engagement… the arguments are sort of normative. You have to do this because you’re working with public money and you have to be accountable or… which is ok, but, I really believe it’s better than that. It’s more important than that. I really believe it’s good to discuss with philosophers, with ethicists, with citizens, with patients… For the quality of your research. To be stretched. It’s another type of checks and balances than the ones which are done in peer review. It doesn’t replace peer review, it’s just another level. To look at the relevance, to, yeah… to be confronted with questions you probably haven’t ever asked… To be better in communicating which… Better communicating will help you better thinking. I mean I think there is a lot of quality gains*.*” (FA)*

But although some perceived science communication to be an essential component of quality work, others saw it as a component which did not indicate the quality nor the efforts invested in the research. Some researchers even supported that the quality of science might be threatened by the lack of quality control of social media.

> *“I feel there is also a kind of danger in those things, because for example I follow some researchers on Twitter, which have a very… I feel that they’re on Twitter all day long I’d say, and everybody follows them… But it’s not… the research is not always that good, but because of the fact that everybody is following, this is going to be the new reality, and I start to… yeah… These things, worldwide, have <an> impact on the research impurities, and it’s shifting towards…. yeah, it’s not controlled, the quality of those things*.*” (Researcher)*

This perspective was echoed by a participant from the RIO group, who admitted having faced substantial resistance from researchers when presenting an action plan meant to promote and value science communication in her institution. This RIO received responses such as “*yeah… that’s the one who’s always with his head in the newspapers, but is he writing A1s*^*2*^*?*”, and concluded that researchers might not be in favour of such a shift. In sum, even though science communication is an important aspect currently put forward in new evaluation processes and policies, researchers do not all agree on its value and its impact on the quality of science.

##### Openness

Openness also raised diverse thoughts from our interviewees. Although most agreed that open science and transparency were important or even “*necessary for the community of researchers”* (PMI), some doubted that open science would help foster integrity, proposing that it might simply bring a “*different level of cheating*” (RIL). We also understood from researchers and especially from research students that the fear of ‘being scooped’ was still too vivid for openness to fully happen, at least before publication. PhD students expressed frustration but also helplessness towards their will to be more open, conceding that the risks of losing their data tended to overcome their will to be more open.

> *“P1: Yeah we are now trying, or in our group someone is trying to put up a database for all of the data on [our topic]. But then researchers would need to hand over their data to make it accessible. And there is a lot of discussion about it, if people would be willing to do that, to hand out your unpublished data… I think it will help the research, and it will help patients, but I don’t know if everyone is willing, I don’t know if I would be willing to, just put it in…*
>
> Interviewer: Would you all… would you be willing to put your data in a server?
>
> *P2: I had the question once, but by a supervisor, and we’re PhD students so you asked, he said ‘No, no we’re just going to publish first and then when we did that then we can say here’s the data’.*
>
> *[…]*
>
> *P3: The problem with research is also it’s really a competition in research. I also have it now that I can’t present on a congress because there are only three articles published on the subject I’m studying, so the supervisors are scared if I make a poster or I present that other researchers will get interested in the same topic, and then, if they publish first all I’m doing is a waste of time… not exactly waste, but… yeah… so I think in research you really have a lot of competition because some people are focusing on the same subject and the data is not published so they will be first, and they want to be first, and… that’s a problem with research. And I think it’s also a problem that no one wants to share their unpublished data because they are scared that someone else will go and take the data and will publish first and then, you don’t have it anymore*.
>
> *P4: Yeah I completely understand the feeling because what we are doing it’s also new so it’s never been done and my promotor is always so reluctant to let me go and show the data to other people. […] he is always so scared that other people are going to steal his ideas… Sometimes I do understand, but sometimes I’m also like, I don’t really like this kind of environment, it struggles with my personality a lot, I think*.*” (PhD students)*

Beyond open data, issues surrounding open access were also brought up in our interviews. We noticed that PhD students, who are directly affected by the inability to access research article, strongly supported open access. Some university leaders also criticized the monopole of big publishers, sustaining that we faced a growing problem where subscriptions may become “unpayable in the long run” (RIL). Other university leaders however, believed that the model of open access was biased, and sustained that they would not advise their researchers to publish in open access journals, also for financial reasons.

> *“That’s another big issue. That’s the open access eh? The model of the open access is unfair because the journal makes the profit by publishing because the author has to pay. So I think the review process is probably more biased. […] I believe. I think that… OK, there is, in my very small field, there is some open access journals, and I feel like whatever review you do they all get published because they get the money. So, is that a good solution? No I don’t believe it’s a good solution*.
>
> Interviewer: Yeah. So would you not advise to your researchers to…
>
> *Participant: I don’t, no, because we don’t have the money (laughs)” (RIL)*

One editor or publisher explained that bad publicity surrounding open access journals may come from the unfortunate reality that the open access model “*opened the door for a number of the so called predatory journals*” (EP). Nonetheless, this interviewee also sustained that “*in the end of the day* [journal quality] *depends on the editorial process of the journal and on the editorial criteria of the journal*” rather than on the publication model.

A number of other indicators raised polarized views, such as the need for societal benefit, and the need for focused areas of expertise. In sum, respondents agreed that current research evaluations were sub-optimal. Nonetheless, disagreements on the specific indicators which should and should not be used in attributing success suggest that solutions are far from simple.

#### A wish for change…

At the end of our interviews, we give participants a ‘fairy wish’ and ask them to describe one change they believe needs priority in science. Although not all answers targeted research assessments, the majority of respondents discussed changes relating to research assessments or research funding as their ‘fairy wish’. In changing research assessments, the need to value quality over quantity, to reduce output pressure and competition, and to broaden and adapt indicators of success to reflect not only the output, but other aspects of science were mentioned. In changing research funding, the need for fairer evaluations and distribution, including the suggestion that resource are not distributed based on assessment but rather equally distributed among scientists, and the wish for long-term funding schemes and baseline research allowances were ‘wished’ from our participants. Appendix 6 illustrates these ideas with a selection of quotes from diverse participants.

### Optimizing research assessments

Despite persisting disagreement on the content of good research assessments, several respondents proposed concrete recommendations on the form research assessments should take. Four main characteristics were put forward as essential for fair and representative evaluations (See Table 2 for sample quotes representing the four criteria).

**Table 2.**
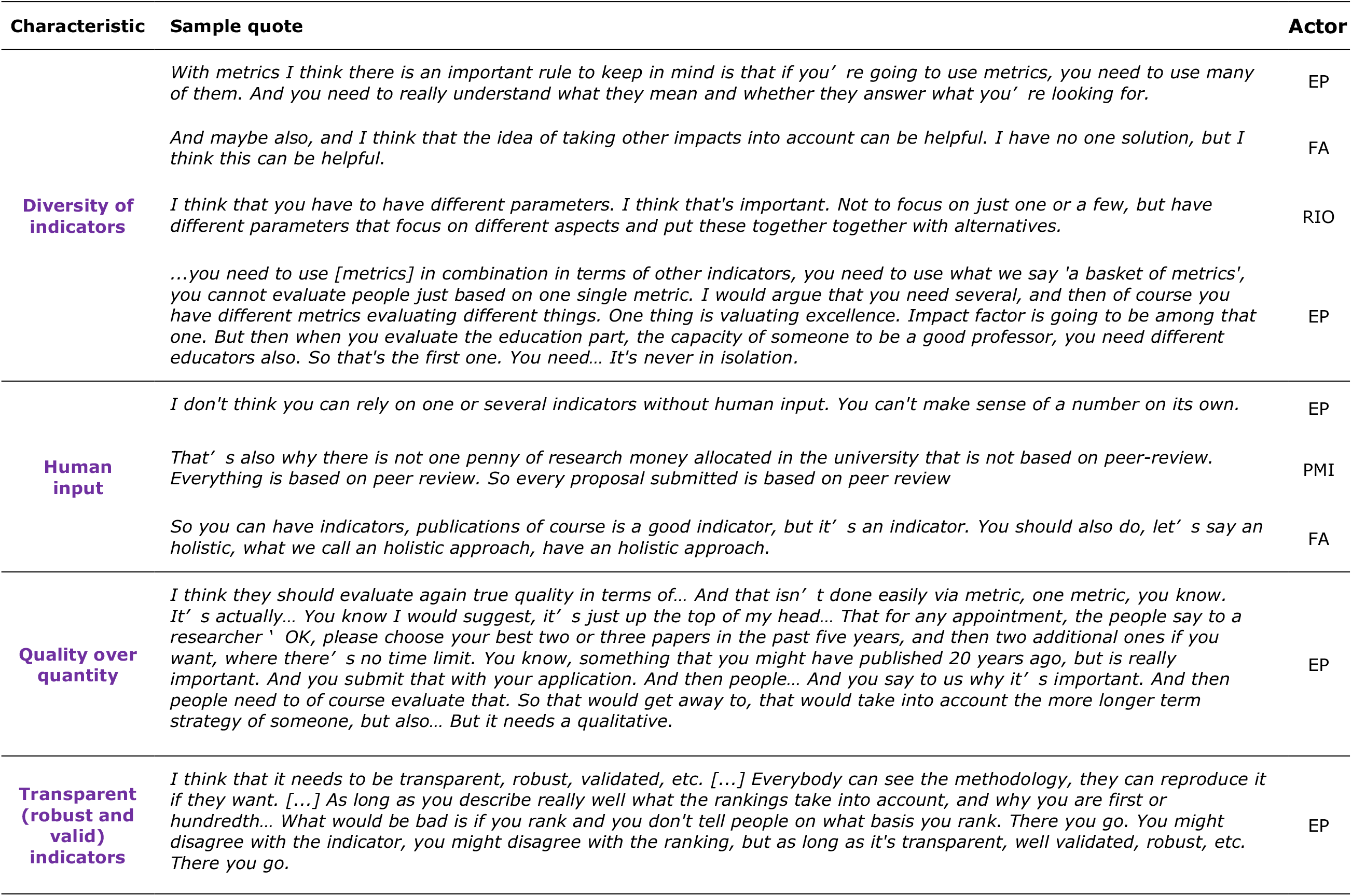
Characteristics for good research assessments.

#### Diversity of indicators

First, many interviewees mentioned that it is essential to use a diversity of indicators to be able to measure different aspects of research. Many respondents worried about the current overreliance on outputs (i.e., especially publications and impact factors). Interviewees sustained that relying on one or few metrics generated important biases, opened the door for manipulation, and ignored important processes which relied both on different metrics and on other type of evaluations, such as openness, societal impact, or science communication

#### Human input

Second, respondents also sustained that it was necessary to have *human input* — in the form of peer review — in the evaluation process to capture what some called a holistic view of success. Peer review was, however, said to also share important weaknesses which must be taken into account. Among those, (i) the *potential for conflicting interest* (especially worrisome to researchers and students who perceived that funding depended more on status and network than on the quality of the project proposed), (ii) *conservatism* (an issue we will explore further in the associated paper, (9)), (iii) *subjectivity*, and (iv) *costs*^*3*^ were mentioned. One research funder proposed that repeating evaluations in different contexts, institutions, and with boards of mixed affiliations could help balance these problems. Another respondent proposed that, to reduce the costs and increase the availability of peer-review, peer-review itself should be rewarded in research assessment.

> *Why shouldn’t people be given credit for doing this kind of work? It’s really important work, it keeps the whole academic system alive. So I think it’s crazy that it’s not included as a, you know, a metric, a possible metric or an indicator of being a successful scientist! (RIN)*

#### Quality over quantity

Third, the importance of evaluating the quality over the quantity was raised many times by many different research actors. Many sustained that presenting only a subset of the most relevant work (e.g., three papers most important to the researchers, and why) could help by permitting in depth evaluation rather than reliance on quantity and metrics. Nevertheless, funders and policy makers both mentioned that despite criticism from researchers about the over reliance on quantity, peer reviewers — generally researchers themselves — often asked for the full list of publications, the H index, or other quantifiable indicators when evaluating proposals, even when the proposal was purposively adapted to contain only a subset of relevant work. Overcoming this quantifiable culture thus seems to be a must for initiating a change.

#### Transparent, robust, and valid indicators

Finally, the transparency, robustness (consistency between evaluations) and validity (measuring what is intended) of indicators were also mentioned as a requirement for good evaluation. These last criteria are basic criteria for any reliable metric, yet they are not always met by newly proposed indicators, and the way current indicators are used sometimes compromises the validity of the intended measure (e.g., assessing *quality* of *single* publications using the impact factors, which qualifies average *journal* citations). Added to these four essential characteristics, the importance of being consistent in how evaluations are conducted while considering differences in fields and disciplines were often raised by interviewees.

### Short summary of findings

Our investigation of the perspectives of success in science reveals that the way in which we currently define science and the way in which we assess scientific excellence generates conflicting perspectives within and between actors.

First, we realised that the way in which researchers define their personal successes was not necessarily standard, and that definitions of successes seem to change with different contexts, demands, and career stages. For instance, the desire to make a change in society was particularly strong in early career researchers, while more established researchers also valued simple curiosity, and relational successes.

When involving all different research actors, we were able to build a representation of success which was nuanced and multifactorial. Success appeared to be an interaction between characteristics from the researcher (Who), research outputs (What), processes (How), and luck. Interviewees noted that current research assessments tended to value outputs but to largely ignore processes, even though these were deemed essential not only for the quality of science, but for the collegiality and the sense of community that unites scientists. Luck was thought to play a crucial role in success and was often used to explain cases where evaluations of success were considered unfair: bad luck explained the lack of reward for excellent researchers, while good luck explained that regular researchers moved ahead without deserving it more than others.

Interviewees generally agreed that current research assessments did not capture the whole picture, and there were a number of disagreement on the specific indicators used to attribute success. The relevance of focalising on publications, impact factors, science communication, and openness in research assessments raised strong and opposing views. When asked what they would change in science, a majority of respondents targeted the way in which research is being assessed and rewarded, sustaining that there is an urgent need for fairer distribution of resources and rewards in science.

Finally, interviewees provided insights on the characteristics they considered essential to any fair and representative assessments. Among those, interviewees sustained that we need a diversity of indicators to allow a balanced and diverse view of success; that assessments should not blindly depend on metrics but also need human input; that we must value quality over quantity; and that any indicators used must be transparent, robust, and valid.

## DISCUSSION

To advance or even maintain their career, researchers need to be successful. But meanings of success in science are not univocal. Different research actors shared their perspective with us, depicting success as a multi-factorial, context-dependent, and mutable, factor which is difficult to define. Unsurprisingly, translating the complex idea of success into concrete assessments is challenging. In many cases, our participants’ personal views on success conflicted with how success is currently determined by research assessments. We are not the first to showcase this conflict: research assessments have been under the radar for quite some time and the discussion on how to improve them continues to grow. The current paper adds to this discussion by showing that, even when considering the perspectives of different research actors, research assessments generate a lot of criticism. One recurrent criticism was the fact that current assessments over-rely on research *outputs*, thereby ignoring, if not discouraging, important *processes* that contribute to the quality of research. This issue is central to the current discussions on the topic. For instance, just a few months ago, Jeremy Farrar, director of Wellcome UK, sustained that the “relentless drive for research excellence has created a culture in modern science that cares exclusively about what is achieved and not about how it is achieved” (15). Resonating this perspective, The Hong Kong Principles for Assessing Researchers sustain that researchers should be assessed on the process of science, including responsible practices (principle 1), transparency (principle 2), and openness (principle 3), as well as a diversity of research activities, scholarship, and outreach (principles 4 and 5). Part of the criticism when assessing outputs also comes from the dominance of inflexible and reductionistic metrics, an issue that was also significant in our findings. Echoing such concerns, the Declaration on research assessments (DORA; 2) directly advocates against using the impact factor for individual evaluations, while the Leiden Manifesto and the Metrics Tide reports pledge for the development and adoption of better, fairer, and more responsible metrics (3, 4). In this regard, our interviewees raised issues which are at the heart of current discussions on research assessments. Yet, our findings also reveal that perspectives on specific assessments remain multi-sided, and that priorities and desired changes are far from univocal. In connecting these different perspectives, we realised that a key question often remains unanswered in the debate on research assessments, namely, ‘What do we want from research assessments?’

Considering both the views on success and the perspective of the problems that were raised in our project (9), we identified three main objectives of research assessments.

### Researchers

First, one of the objective of research assessments appears to promote and value good researchers. Our respondents suggest that success in science has an important individual facet. Assessments were often described as a way to reflect personal merits and to provide recognition for skills, competencies, and efforts. Research assessments are thus expected to be, at least in part, meant for the fair recognition of researchers’ accomplishments. And indeed, fairness was central to our discussions on success. Interviewees expressed their concern for fairness by blaming luck (and bad luck) for inexplicable successes (or lack thereof) and by worrying that connections, seniority, and renown could yield unfair advantages which are not related to genuine merit (9). Such concerns suggest that current research assessments fall short on many points if their main objective is to fairly reward researchers.

But valuing researchers is not only a matter of fairness. Valuing researchers also means building capacities and nurturing autonomy in order to create strong and sustainable research units. Accordingly, if the goal for research assessments is to promote excellent researchers, they should also facilitate, support, and sustain the creation of strong research teams. This perspective suggests that we should not only recognise personal merit, but also teamwork, diversity, inclusion, and collegiality. Yet, our respondents identified important problems in current research climates which may inhibit these essential features and foster competition, mutual blame, and mistrust instead (9). Many of these problems have been echoed in past research, such as the precariousness of research careers (16), the vulnerability of researchers’ well-being (e.g., 17, 18), and the perceived lack of institutional support for researchers (19). Knowing that researchers’ perceptions of research climates can directly influence research practices (20, 21), it seems urgent to address issues embedded in research climates before assessments can fulfil the advancement of strong, sustainable, and flourishing research teams.

### Society

Other approaches rather focus on the benefits that science can bring to society. A common argument for the need to benefit society is the fact that science is primarily financed through public money and should thus profit back (tangibly and intellectually) to society. Following this perspective, research assessments should aim to ensure that scientists involve, communicate, and implement their findings within society. Applicability of research findings, public engagement, science communication, open access, and feasibility would be at the heart of this objective. But in practice, research assessments often neglect societal benefits (22). Our interviewees offered polarised views on this topic, with some valuing and others downgrading science communication. These polarised views suggest that the neglect for societal benefit may be deeply rooted within cultures. We have also explained that, at least within our modest sample, the values of open science and the desire for implementation seemed to diminish as career advanced. While this finding may be anecdotal, it could also suggest that the broad neglect for societal benefit in current assessments shapes researchers’ perspectives of success, encouraging them to prioritize competition and metrics over openness and societal value. Consequently, if research assessments aim to promote and value societal benefit, they might first need to reconsider research cultures.

### Science

Finally, we should not overlook research’s primary and inherent goal of advancing science and knowledge. Knowledge is often described as the common objective and the end in itself of science. In this regard, research assessments should ultimately encourage the advancement of science. Two aspects are then essential to consider here. First, to advance science, we need to ensure that research is conducted with integrity. Assessments should thus encourage the *processes* which maximise the integrity and the quality of research. Yet, openness, reproducibility, rigorousness, and transparency were recurrently mentioned as missing from current research assessments. Certain aspects of current assessments were even thought to discourage integrity and research quality. Many interviewees supported that the lack of consideration for negative results caused tremendous research waste, that competitiveness of assessments compromised collaborative efforts and transparency, and that the current focus on ‘extraordinary findings’ discouraged openness and transparency (9). Evidently, if the reason for assessing research is to promote the advancement of science, processes which foster integrity must be given due recognition. But even when integrity and quality of research are ensured, advancing science requires continued innovation, creativity, and productivity. According to our findings, this is where most research assessment currently focus. Publications counts and impact metrics ensure that ‘new’ knowledge is created, and that the pace of creation is fast enough. Yet, the overemphasis on outputs and the negligence of processes was highly criticized by our interviewees. Current assessments, for instance, were said to shift researchers’ focus from ‘what is needed to advance the field’ to ‘what is sexy to publish’, or ‘what will attract funding’. Current assessment systems were further criticized for their conservatism and for the difficulty to pass disrupting and truly innovative ideas through peer-review (9). Short-term funding schemes and the high pace of research evaluations were also criticized, with interviewees sustaining that innovative research requires long-term investments and sufficient freedom for failure — two conditions which were noted as missing in current funding schemes. In sum, both the overlook of research processes and the expectation of quick, positive research results suggest that current research assessments are not optimized to help advancing science, and that they need to be addressed.

Our findings do not provide an answer to what research assessments *should* aim for, but rather illustrate that resolving this first — and often overlooked — question is already challenging. The variety of answers we collected suggest that the perceived objectives of research assessments may differ from person to person, and that specific actor roles may come into play. In theory, it seems reasonable that universities aim to create sustainable and empowered research teams, that funders and policy makers aim to maximize the societal value of science, that publishers, editors and researchers aim to contribute solid advances to the existing pool of knowledge, etc. But such simple perspectives do not reflected reality, where research actors are themselves individuals with personal perspectives, experiences, and convictions. The complex association of perspective indubitably provides richness to the scientific system, and it would be absurd, if not damaging, to aim for a single unified perspective. Yet, the ‘end’ goal of research assessments is often obscured from discussions on the topic. Most discussions aim to find the ‘means’ (i.e., metrics, indicators) to fit the ‘end’, but fail to define the end of research assessments. Ensuring that the discussion on research assessments listens to the perspectives of all research actors — including the forgotten voices such as early career researchers — and that all parties are transparent and explicit about what they wish to achieve by assessing researchers may be a first step for an open dialogue to enable concrete changes to take place.

## CONCLUSION

The present paper describes perspectives from different research actors on what defines and determines success in research. In their answers, interviewees raised a number of shortcomings about the approaches currently used for assessing success in science, and these shortcomings lead to important problems in the functioning of science (see the associated paper 9). Most notably, participants sustained that current research assessments place too much emphasis on research outputs and on quantity, while they largely overlook research processes and indicators of quality.

Issues with research assessments have been on the priority agenda for some years already. But although reflections and ideas for change are on the rise, concrete changes are still moderate and sporadic. In this paper, we bring the debate one step back to ask ‘What do we really want from research assessments?’. Are assessments meant to value and encourage good researchers, to benefit society, or to advance science? We sustain that current research assessments fall short on each of these core objectives, and need to be addressed.

Assessing researchers is an issue that has high stakes, not only for individual researchers who wish to continue their career and seek recognition, but also for the future of science. Our combined papers (9) reiterate that current research assessments need to be revisited, that all research actors must be involved in the discussion, and that the dialogue must be open, inclusive, transparent, and explicit. Acceptability, trust, and joint efforts can only be increased if all actors are involved, understand the other’s perspective, and work together to build a solution.

## LIST OF ABBREVIATIONS

COREQ: COnsolidated criteria for REporting Qualitative research checklist
DORA: San Francisco Declaration on Research Assessment
EP: Editor(s) or publisher(s)
ESF: European Science Foundation
EUA: European Universities Association
FA: Funding Agency(s)
LT: Laboratory technician(s)
PMI: Policy Maker(s) or influencer(s)
PostDoc: Post-Doctoral Researcher(s)
QUAGOL: Qualitative Analysis Guide of Leuven
RCC: Researcher(s) who changed career
Re-SInC: Rethinking Success, Integrity, and Culture in Science
RIL: Research institution leader(s)
RIN: Research integrity network member(s)
RIO: Research integrity office member(s)
WCRI: World Conference on Research Integrity

## Acknowledgements

The authors wish to thank Raymond De Vries, who substantially contributed to the Conceptualization, Methodology, Resources, and Validation of the present project. The authors also wish to thank Melissa S. Anderson and Brian C. Martinson and Raymond De Vries for sharing their focus group guides which constituted the foundation of ours (Resources). We also wish to thank Ines Steffens, Inge Thijs, and Igna Rutten who were essential in helping us organise focus groups and recruit participants (Resources).Finally, and most importantly, we want to thank all those who participated in our interviews and focus groups. We know that we forced ourselves in the very busy schedules of many a participant, and we are sincerely grateful for the time, efforts, and precious thoughts that participants generously shared with us.

## DECLARATIONS

### Ethics approval and consent to participate

The project was approved by the Medical Ethics Committee of the Faculty of Medicine and Life Sciences of Hasselt University (protocol number CME2016/679), and all participants provided written consent for participation.

### Consent for publication

Participants provided written consent for participation and for publication and dissemination of the findings from this project.

### Availability of data and material

#### Data

Given the sensibility and risk of identification of qualitative data, we are unable to share full transcripts. In the manuscript however, we attempted to be as transparent as possible by providing numerous quotes in the text and in tables to exemplify and support our claims.

#### Materials

Our focus group and interview guides, as well as the consent forms and information sheet we shared with participants are available on the registration of this project (11).

## Authors’ contributions

Authorship contributions: NAB contributed to the Conceptualization, Project administration, Methodology, Resources, Investigation, Data curation, Formal Analysis, Visualization, Validation, Writing – original draft, and Writing – review & editing. WP contributed to the Conceptualization, Funding acquisition, Methodology, Resources, Validation, Supervision, and Writing – review & editing, both intermediate and final versions.

## Authors’ information

**NAB** is a PhD student supervised by WP at the faculty of Health and Life Sciences in Hasselt University, Belgium. Her research interests are predominantly in research integrity and publication ethics, especially targeting research assessments and the way in which scientific systems influence the integrity and the quality of science.

**WP** is associate professor at the faculty of Health and Life Sciences in Hasselt University, Belgium. As a medical ethicist with expertise in research ethics, he has increasingly devoted attention to education and research in the field of research integrity, with a specific interest in research culture and the definition of success in science. He is supervisor of NAB.

## Footnotes

1 Initially, we were tempted to classify these four categories in *Products* of success (the What) and *Potential* for achieving success (the Who, How, and luck). However, after several discussions, we realized that doing so may reinforce the perspective that products are the ones which truly indicate success, while potentials are simply increasing the chance of yielding better products. As we will describe later on, many of our interviewees considered the ‘Who’ and especially the ‘How’ to be genuine successes in themselves. In this regard, we intentionally kept the four categories together as each representing successes in themselves.

2 A1’s are a category of publications in research assessments in Flanders. They relate to articles included in Web of Science’s Science Citation Index Expanded, Social Science Citation Index and/or Arts and Humanities Citation Index, whose document type is labelled as “Article”, “Review”, “Letter”, “Note” and/or “Proceedings Paper”; or in journals included in the Journal Citation Reports of Web of Science.

3 This aspect was raised when discussing external expert panels for integrity issues, in which one RIO mentioned that “They’re paid for leading the report and making the preparations, you have to bring them to your university, you have to put them in a nice hotel, obviously, you have to dine, I mean, the amount of money is just enormous. And then you have… ok what is coming out of this? You have some remarks… […] Yeah I’m a bit critical towards that system”.

