## Appendix 1 for "Rethinking success, integrity, and culture in research (part 1) — A multi-actor qualitative study on success in science"

### RESEARCH TEAM AND REFLEXIVITY

In accordance with the COnsolidated criteria for REporting Qualitative research checklist (COREQ; Appendix 2), and in respect of transparency, we found important to provide further characteristics about the setting and the interviewer at the time of the study.

Besides one early interview with an institution leader in which WP, assistant professor, attended to provide feedback about the interview, all other interviews and focus groups were conducted by NAB, with no additional non-participant or assistant.

NAB is a female PhD student in the Faculty of medicine and life science of Hasselt University, Belgium, with a background in cognitive neuroscience and bioethics. Coming from Canada, NAB had the advantage of bringing a certain neutrality in the interviews by not being strongly affiliated with one or another Flemish region, and by not corresponding to an established research group.

Before conducting the interviews and focus, NAB followed courses about developing interview questions, conducting focus groups, and analysing qualitative data offered from Flemish universities and from the Flanders' Training Network for Methodology and Statistics (FLAMES). In addition, she used the resource books from the Focus Group Kit by Richard A. Krueger and David L. Morgan (Morgan & Krueger, 1998), and discussed with co-author RDV — expert in qualitative inquiries and part of the team that built the original guide upon which we inspired ours — to gain insight on building, conducting, and analysing focus groups and interviews.

Besides a few exceptions, NAB had no prior relationship with most participants, and the first contacts were established with the invitation email. No repeat interviews were carried out. Before the interview, NAB described the project briefly and explained the purpose of the interview informally. On some occasions where interviewees were anxious to know more about the project in advance, NAB would email the main themes targeted, but would not share the interview guide with participants by fear that this may lead to rote answers.

#### Bias and assumptions

NAB holds the view that research integrity is largely determined by the research system, and the interview guide was necessarily not unbiased to this perspective. Nonetheless, if participants shared a different view (e.g., if they believed that integrity was solely a matter of personality), NAB was careful not to contradict or bias interviewees' ideas towards her perspective. In re-reading quotes with the research team, we were careful for possible misinterpretations, and when quotes were interpreted differently by WP or RDV, we adapted the nodes and interpretations to make sure they fit the words of the participants.

#### Study design and interview/focus group setting

Interviews and focus groups were conducted in private meeting rooms or offices or, according to preference, in public spaces (N=2) or through video call (N=3). One of the interview conducted through video call had some sound and connection problems, but the other video calls went very smoothly.

Interviews lasted on average 60 minutes, depending on the time granted by the interviewee (range from 34 to 80 minutes). Focus groups lasted around 120 minutes each and included a five-minute break.

All interviews were audio recorded and transcribed verbatim by the interviewer (NAB) or a university-approved transcription service. Transcripts were not returned to participants except in select cases where participants expressed a wish to monitor their answers, and in cases where the quotes of interest might have jeopardized the confidentiality of participants. No repeat interviews were undertaken. After most interviews, the interviewer filled a self-questionnaire about the interview to note any abnormalities and general feelings of the interview data.
