## Appendix 2 for "Rethinking success, integrity, and culture in research (part 1) — A multi-actor qualitative study on success in science"

### COREQ CHECKLIST (COnsolidated criteria for REporting Qualitative research)

| Topic | Item No. | Guide Questions/Description | Reported on Page No. |
|---|---|---|---|
| **Domain 1: Research team and reflexivity** | | | |
| *Personal characteristics* | | | |
| Interviewer/facilitator | 1 | Which author/s conducted the interview or focus group? | Appendix 1**Error! Bookmark not defined.** |
| Credentials | 2 | What were the researcher's credentials? E.g. PhD, MD | Appendix 1 |
| Occupation | 3 | What was their occupation at the time of the study? | Appendix 1 |
| Gender | 4 | Was the researcher male or female? | Appendix 1 |
| Experience and training | 5 | What experience or training did the researcher have? | Appendix 1 |
| *Relationship with participants* | | | |
| Relationship established | 6 | Was a relationship established prior to study commencement? | Appendix 1 |
| Participant knowledge of the interviewer | 7 | What did the participants know about the researcher? e.g. personal goals, reasons for doing the research | Appendix 1 |
| Interviewer characteristics | 8 | What characteristics were reported about the inter viewer/facilitator? e.g. Bias, assumptions, reasons and interests in the research topic | Appendix 1 |
| **Domain 2: Study design** | | | |
| *Theoretical framework* | | | |
| Methodological orientation and theory | 9 | What methodological orientation was stated to underpin the study? e.g. grounded theory, discourse analysis, ethnography, phenomenology, content analysis | Page 8-9 |
| *Participant selection* | | | |
| Sampling | 10 | How were participants selected? e.g. purposive, convenience, consecutive, snowball | Page 7 |
| Method of approach | 11 | How were participants approached? e.g. face-to-face, telephone, mail, email | Appendix 1 |
| Sample size | 12 | How many participants were in the study? | Page 6 |
| Non-participation | 13 | How many people refused to participate or dropped out? Reasons? | — |
| *Setting* | | | |
| Setting of data collection | 14 | Where was the data collected? e.g. home, clinic, workplace | Appendix 1 |
| Presence of non-participants | 15 | Was anyone else present besides the participants and researchers? | Appendix 1 |
| Description of sample | 16 | What are the important characteristics of the sample? e.g. demographic data, date | Page 6 |
| *Data collection* | | | |

| | | | |
|---|---|---|---|
| Interview guide | 17 | Were questions, prompts, guides provided by the authors? Was it pilot tested? | Appendix 1 |
| Repeat interviews | 18 | Were repeat inter views carried out? If yes, how many? | Appendix 1 |
| Audio/visual recording | 19 | Did the research use audio or visual recording to collect the data? | Appendix 1 |
| Field notes | 20 | Were field notes made during and/or after the inter view or focus group? | Appendix 1 |
| Duration | 21 | What was the duration of the inter views or focus group? | Appendix 1 |
| Data saturation | 22 | Was data saturation discussed? | — |
| Transcripts returned | 23 | Were transcripts returned to participants for comment and/or correction? | Appendix 1 |
| **Domain 3: analysis and findings** | | | |
| *Data analysis* | | | |
| Number of data coders | 24 | How many data coders coded the data? | Page 8-9 |
| Description of the coding tree | 25 | Did authors provide a description of the coding tree? | Page 13 and (1) page 13 |
| Derivation of themes | 26 | Were themes identified in advance or derived from the data? | Page 8-9 |
| Software | 27 | What software, if applicable, was used to manage the data? | Page 8-9 |
| Participant checking | 28 | Did participants provide feedback on the findings? | Appendix 1 |
| *Reporting* | | | |
| Quotations presented | 29 | Were participant quotations presented to illustrate the themes/findings? Was each quotation identified? e.g. participant number | Throughout |
| Data and findings consistent | 30 | Was there consistency between the data presented and the findings? | Throughout |
| Clarity of major themes | 31 | Were major themes clearly presented in the findings? | Page 13 and (1) page 13 |
| Clarity of minor themes | 32 | Is there a description of diverse cases or discussion of minor themes? | Throughout |

1. Aubert Bonn N, Pinxten W. Rethinking success, integrity, and culture in research (Part 2) - A multi-actor qualitative study on problems of science. bioRxiv. 2020.
