## Appendix 3 for "Rethinking success, integrity, and culture in research (part 1) — A multi-actor qualitative study on success in science"

### General interview guide

*A part of my research is to explore the views of different actors that contribute to the research system.*

*To protect your privacy, I want to avoid disclosing your specific job title and to place you in one of bigger category of research actors. I may use a higher level of details to describe the type of participants included in each category, but I won't link direct quotes with company or institution names.*

*I placed you in the category ***actor group***. Does that sound good to you?*

#### Introduction and information on respondent's career

1. Before anything, I would like you to **describe your work** to me, in your own words.

   *Prompt: In* a broader perspective, what would you say is your **role is in the scientific system**?

2. In this job, you obviously **care for scientific excellence**. How would you say you fulfil this goal in your work?

   *We will get back to this a bit later. For now, I will change topic and I want us to talk about success as this is an important topic that we are trying to understand in the project.*

#### Success in science

3. First, try to think about scientists you've known that were very successful. What do you think made these scientists **successful**?

   Prompt: Which characteristics do you think are most important to advance a researcher's career?

4. Do you feel like these characteristics are **captured in current research assessments** and evaluations? In which ways?
5. (If time allows) What do you feel that your actor group **should do to promote successful science**? Do you see that happening?

#### Tensions or conflict between success and integrity

6. You mentioned that X, Y, Z are criterions that indicate success in research. Do you think that these are **also indicators of quality**? Sound research?

   *Prompt: Which criterions do you think **indicate the quality** of the research?*
   *Prompt: Which criterions do you feel are **not suited** to indicate the quality of the research? Explain.*

7. Does it happen that you see excellent researchers but for some reason these researchers **don't succeed in getting ahead** with their career? Can you give me some examples?
8. Do you feel that the way in which success is attributed allows to for **emerging scientists** to become successful?

### Current problems

Let's change the topic now; leave aside success for a bit and look at when science is not at its best. Like I said, I am not here to denounce or condemn cases, so I will make sure to protect the confidentiality of cases you may discuss.

9. Have you ever had to **deal with science** which you considered was **not really in line with the rules of science**? What happened?
10. Can you give me precise examples of the elements that you consider **signs of bad or sloppy research**? What are **red flags**?

### Motives for bad practices

11. **Why** do you think bad research practice happens?
12. Do you think **anyone could** end up in such a situation or only types of people?

### Responsibilities towards integrity

We have already discussed how to promote successful science, now I would like to gather your thoughts on how to prevent sloppy research.

13. What do you think should be done to **prevent bad science** from happening?
14. **Who** should take the lead to make these changes happen? Who else should be involved?
15. What do you consider is the **responsibility of your *actor group*** to protect integrity?
16. Where does your responsibility **end**?

### One change

Finally, if *you could **pick one important change** that needs priority right now in how research works, what would it be? How do you think this change could be done?*

### (if time allows) Alternatives

17. If there were no rules for evaluating scientists, and you could **start from scratch**, what would you like to look at when assessing scientists?

    *Prompt: What are the characteristics that **YOU think are most important** for researchers to do good research?*
