## Appendix 4 for "Rethinking success, integrity, and culture in research (part 1) — A multi-actor qualitative study on success in science"

### GENERAL FOCUS GROUP GUIDE

Intro:

1. Who you are
2. What is your area of research
3. Describe a typical day of work
4. What's your favourite ice cream flavor

#### Scientific career

Before starting, I would like to know a little more about your career as a researcher.

1. Specifically, I would like to know what is it that makes your work so great? What do you feel is most satisfying, most rewarding about your career?

   *Prompt: When people ask you why you chose to be a researcher, what first comes to mind?*

We will get back to this a bit later. But for now I will change topic and I want us to talk about success.

#### Success in science

2. Think of a person in your field who you think is **very successful**. (It doesn't have to be one person in particular, it can just be some characteristics of many different people, can be yourself in 20 years…) How do researchers **become successful**? What, in your view makes this person a success?

   *Prompt: What are the most important factors for advancing in your career?*
   *Prompt: What are the funders and the employers looking at?*

3. Now imagine that I am a newcomer in your field and I ask you what I must do to stay on the top, **what would you tell me**?

So you say that successful scientists are generally scientists who do X, Y, Z.

4. Do these successful scientists **reflect or mirror the kind of scientist you want to be**? Do you have such aspirations for success?

#### Tensions or conflict between success and integrity

5. As we discussed, you point out that funders and employers look at X, Y, Z… Do you think that **these criterions for success indicate outstanding or excellent research** (e.g., appropriate methods, relevant topic, high quality work)?

   *Prompt: Which criterions do you think **indicate the quality** of the research?*
   *Prompt: Which criterions do you feel are **not suited to indicate the quality** of the research? Explain.*

6. Now try to think of a **colleague** who, in your opinion, **does good research but cannot reach success in science**?

   Prompt: *What in your opinion explains that this researcher cannot reach a successful career?*

7. **What would you say to this researcher to help him/her get ahead**?

### Current problems

Let's change back the topic now; leave success aside for a bit and discuss what it is like to be a researcher. So you remember that at the beginning of the discussion, I asked you about the aspects of research that make you like your career. Now I want us to talk about the other side of things, about what **frustrates** you as a researcher.

8. So let's say I am a **newcomer** in your field. I just started working in your lab and I am not sure whether I should follow a scientific career. If I asked you what are the **most frustrating things about working in science**, what would you tell me?

   *Prompt: what would you tell me are some of the biggest frustration I could encounter?*

9. All right, so as we have discussed, being a researcher is not necessarily always easy. It can sometimes happen that things go really wrong. Have you **ever seen or heard** of a situation in which you thought research was conducted in a way that was **against the 'rules' of science'**? What happened? What did/would you do?

### Motives for bad practices

10. **Why** do you think researchers were acting in this way?
11. Do you think **any researcher could end up** in such a situation?

### CURRENT VIEWS ON RESPONSIBILITY

12. **What** do you think should be done to **prevent bad science** from happening?
13. **Who should take the lead** to make these changes happen? Who else should be involved?
14. **What can you do**?

    *Prompt: Do you feel like you miss something to be able to change things yourself?*

### Solutions

*To finish, I would like to ask a more concrete question.*

15. Finally, if you could pick **one important change that needs priority** right now in the research system, in how science works, what would it be?

    Prompt: How do you think this change could be done?

### Personal success

Now before we finish, I want you to think back about the discussion we have had on success, and on criterions that are most often used to evaluate a research career. But now, I would like you to think about yourself as a researcher, and to think about your strength, about what makes you feel accomplished in your work. **What do you think is your biggest contribution to your work**, or **something that you think is key to be a good researcher**, regardless of the criterions we have said before. (For example, maybe you think that the fact that you brush your teeth after lunch is key to the success of your research team.)

I will not ask you to discuss it this time, but I would like everyone to take one of these little pieces of paper. On the piece of paper, I would like you to make up 3 to 5 criterions for funders and employers. I want you to think about what you consider your biggest contributions in your work, and to make up criterions you would think, if funders and employers evaluated, you would have better chances to succeed.

### Summary

Is there anything you would like to mention that we failed to discuss today?
