## Appendix 5 for "Rethinking success, integrity, and culture in research (part 1) — A multi-actor qualitative study on success in science"

### SAMPLE QUOTES ON PUBLICATIONS

Sample quotes substantiating arguments against and for using publications as a main determinant for scientific success.

| Argument | Sample quote | actor |
|---|---|---|
| **Arguments against using publications as the main determinant of scientific success** | | |
| Reductionist | *It is a very flawed measure of success in a way. I mean… I don't want to give the impression that… of discouraging any of these successes, you know, I mean publishing very important papers in very selective journals is an achievement, that is very clear. But I think that there are other very important contributions to the scientific enterprise which don't necessarily translate into one of these unit of credit of success, which is a first author publication in a very prestigious journal. And I think that currently we collectively, as a community, do not do enough to actually support and reward these kinds of contributions that are very important for the scientific enterprise* | EP |
| Arbitrary | *Yeah but with publications it's sometimes also just having luck…[…] To me it's not always that you're a good researcher.* | LT |
| | *It is wrong to think that... […] having more publications, it means you're better and better and better, I think it's a very wrong way of thinking.* | PMI |
| | *I have less and less confidence in publishing with the fact that 'who is going to be the reviewer?' 'Is he biased?' 'Is it the journal?'* | R |
| | *The highest journal [of my field] it's all already fixed before with companies, pharmaceutical companies, who will get published their RCTs, it's already all set in advance...* | PhD |
| Perverse | *They do a lot of experiments just to publish. Just to make an article, because they have to have an article before the four years are done. So they do their experiments in function of an article* | LT |
| | *It's my only drive for some things, that it's just publication.* | Res. |
| **Arguments in favour of using publications as the main determinant of scientific success** | | |
| Representative | *So people say, you know, publications don't matter, but at the end of the day there clearly is a link. If you end up publishing in a good journal, then you probably started off with a very good research question, and you probably are a very good researcher. They are not 100% linked, but I'm sure there is a link there.* | RIL |
| Measurable | *"It's the career, it's the way you get the career, it's the number of publications that will count, the number of promotions of PhD theses will count, but for me that's not the most important. I think a researcher who is not... who is publishing (they need to publish of course) but let's say only two A1 publication, or one publication a year, but in the meantime is contaminating other researchers, helping other researchers and is multidisciplinary... That's more valuable for me as a person. But in the academic world,* ***I cannot value that directly****. I'm not in a position that I can say "You are the very best researcher, so I promote you to full professor from associate professor". Because there we still have the numbers that count. And ok, that's the way it is, and that's the whole issue nowadays with researchers. They really get troubled with these numbers."* | RIL |
| | *"I think it would also be a bit difficult to really value a PhD or the PhD project without publications. Because how do you determine that someone has done their best, but unfortunately didn't get any publications."* | PhD |
| | *"I do believe that you have to have some evidence about the process you have made, and the path that you've walked throughout your doctoral thesis. That's why I find it quite normal that you have to have a certain amount of publications in the procedure… "* | RIO |
| Necessary | *"I think I'm going to be the boring one, but I think it is important to have publications and to also be successful in some research grounds every now and then because I feel like it's my... That's what is expected from me, but that's also how you can make the research... you can keep the research going. I think it's one I see as my duty to publish the results, to share them so that others can build further on them and you yourself can build further on them."* | Res. |
| | *"If you don't have the publications you're not noticed. And if you're not noticed, your research might be extremely interesting, but if it's not read, if it's not noticed, what's the value."* | RCC |

Note: Researcher is abbreviated to Res.
