## Appendix 6 for "Rethinking success, integrity, and culture in research (part 1) — A multi-actor qualitative study on success in science"

### SAMPLE QUOTES 'WISH FOR CHANGE'

Sample quotes from the 'wish for change' which relate to changes in the ways success is defined and assessed.

| Actor | Sample quote |
|---|---|
| **CHANGES TO RESEARCH ASSESSMENTS** | |
| ***Value quality over quantity*** | |
| RIO | *I will then insist that the money is spent on projects of high quality. Quality of the research.* |
| Res. | (immediate response) *Take the output pressure away! So you can have more room for quality.* |
| PMI | [Participant] *My wish is that scientific outcomes, papers, pieces, news, are assessed by their intrinsic value, intrinsic scholarly value and not by indirect measures as it is the case right now. Journal impact factors, citation index, et cetera, these are all proxies.*<br>[Interviewer] *What would you say is any intrinsic value?*<br>[Participant] *I think open peer review would solve this problem.* |
| ***Reduce output pressure and competition*** | |
| EP | *I would like to see a world where the pressure is off the researchers, you know, not... there are not pressured, in the world that they can do their research without pressure of publishing in high impact journals, and like to see that there is no impact factor anymore at least not in such a way that there is usually considered today. And that to bring more joy in their life, essentially, because I think that they are so stressed out, and they are always chasing some next step in their career advancement, and they forgot that the science is actually fun thing to do, you know, it can be a way of, you know, living a life, not just working as a hamster in a wheel, you know, just yeah, chasing your own tail or something like that.* |
| EP | *(Laughs!) It is a really tough one. Because I don't see… Do you know Merton's model? […] OK. There is tension, there is obvious tension between the kudos, and the whole system that has been put in place where it's… you have to be special. It doesn't fit. It doesn't fit with the kudos! It doesn't fit with the universalism, etc. So I think that that's where something is wrong. I don't have the solution, but that's what needs to be addressed! [...] I would try to solve that tension that exists right there, to be able to go back to the other communalism, to the universalism etc. You know, the kudos itself.* |
| EP | *Change the reward system! (Laughs) Change the reward system. Completely. Because would then allow everyone A) to publish wherever it's really most relevant, it's not linked to the impact factor any longer… You know if people did that, what I said, and this was not relevant, impact factor was not relevant, and it's really truly about what kind of research career have I had and what research have I done, you know, that is really important, and how does this impact in my field. Then I think everything would change. And, yeah, that would be my biggest wish, and I'm working towards that.* |
| ***Broaden and adapt indicators*** | |
| PhD | *Maybe the cumulative impact factor that they just need to do it really field per field, and not faculty per faculty* |
| RIN | *I think it would be broadening of the criteria for recruitment, promotion, funding. I think if we could really get everybody behind that, it would have a huge impact I think.* |
| PhD | *...maybe looking at PhD as a career. Because now you have only one main outcome, the publications, but in a career you have a lot of competencies that are important.* |
| EP | *If I have a magic wand, I think I would want to get rid of the Impact Factor in research assessment. And getting rid of… You know changing this problem that we started this conversation with. Which is that it's only publication in a certain amount… in a small number of very selective journals that is considered a measure of success. So, you know, I would want a magic number that represents all these other things and that's probably, that's completely unrealistic, but I would want at a minimum the research assessment framework to change to move away from that single dominant measure that is being used at the moment. To appreciate different kinds of contributions much more effectively.* |
| **CHANGES TO RESEARCH FUNDING** | |
| ***Fairness in evaluation*** | |
| PhD | *Z: I think that... I feel that there is a bias that certain groups will always get funding, and smaller universities [...] are really struggling to get like an FWO project funded. So maybe there should be some regulations about it. But I don't know how... maybe restricting the number of projects that the group can apply for. I think they already have some regulations for that...*<br>*M: I think so too.*<br>*[...]*<br>*L: Anonymous selection... (laughs)*<br>*Z: That would be maybe the best.*<br>*M: But that is also important, because a lot of professors or researchers knew people that are the judges, and they have like, the privilege, and will get funding. So anonymous would be better.*<br>*Z: But then maybe you lose the advantage of collaboration. You can't say "but we'll be collaborating".* |

***Fairness in distribution***

| | |
|---|---|
| EP | *Yes, my fairy wish would be a change of the grant system, and I'm saying of that it… in my… so say 80 percent of the money might be divided according to the prevailing system, for the proposals and give 80 percent to the best proposals, and then we have a pile of proposals that are rejected, and make it a lottery, for 20 percent.* |
| RCC | *Participant: More freedom. Less bullshit. More... and maybe the money should be just divided equally or something like that, which is also not really realistic because then the amount would probably be so small that you still cannot do anything. But at least then everybody can not do anything, instead of being, you know, when you have this big in house thing and here is the people that get a lot of money and get a lot of stuff done, and here's me [laughs]. At least we all will be...*<br>*Interviewer: Everyone would be fair.*<br>*Participant: Yeah, it's just not fair. That's it, it's not fair. And if you... I can completely understand why big science people don't go to [small university] because you kill your career if you [go there].* |

***Long-term and baseline funding to increase security***

| | |
|---|---|
| RIL | *I think it's the research funding, but I don't... I just know it should change, but I don't have the answer for you. I think a researcher should not have these short term financing situations. I think that's probably the worst perverse incentive you can give a scientist. I think you should have a tenior track where you require that a scientist prooves him or herself, but once you have an established scientist, they should have some sort of basic funding which could be adjusted based on how they perform, but it should not be this 'yes/no' thing on a four year term which is what most grants are. Because I really need to deliver in four years, and that gives me perverse incentives.* |
| RCC | *But maybe it might be interesting to give people different kinds of contracts. To don't give always these short-term contracts, but give people longer term contracts. But I know that there's a discussion. I know a lot of people say 'well I give the best of myself because I have a short-term contract and the edge is on… I don't know whether the edge should be so strong. I don't know whether the competition should be so strong. I don't know whether that's really helpful. If you really want to achieve trust and if you really want to achieve openness to mistakes, people should feel secure enough to do it. And I think one of the answers is 'you will not lose your job'. So… Maybe job security might be an answer. (RCC)* |
| *FA* | *Hmhm. Well exactly what I said form the start. I think that we should have a very close look at the way we are funding institutions for doing their research. I think this is key […] but there are some elements I recognise, and we recognise, that are worth a good discussion. And one of these elements is that indeed apart from competitive funding, which is important because competition, and what we are doing here can make for good quality research, excellent research, and apart from this competitive funding, you also need some sort of basic funding to give people a chance to start and to launch their career as an academic. Also to do some things that are less fashionable, because also research has its fashions, less fashionable, or less appealing to evaluators at the moment, with which you can prove after a while that there is something in it and then you become stronger to an evaluation panel. So I think that reconsidering the way you are funding research institutions is also letting some pressure, or diminishing some pressure on institutions like us. I think you get better competition, by also making it less stringent. Maybe this sounds as a paradox, but I don't think it is.* |
| *Res* | *Start-up money? For creative plans which are not judged from the beginning?* |

Note: Researcher is abbreviated to Res.
